# Genomic resources for a historical collection of cultivated two-row European spring barley genotypes

**DOI:** 10.1101/2023.03.06.531259

**Authors:** Miriam Schreiber, Ronja Wonneberger, Allison M. Haaning, Max Coulter, Joanne Russell, Axel Himmelbach, Anne Fiebig, Gary J. Muehlbauer, Nils Stein, Robbie Waugh

## Abstract

Barley genomic resources are increasing rapidly, with the publication of a barley pangenome as one of the latest developments. Two-row spring barley cultivars are intensely studied as they are the source of high-quality grain for malting and distilling. Here we provide data from a European two-row spring barley population containing 209 different genotypes registered for the UK market between 1830 to 2014. The dataset encompasses RNA-sequencing data from six different tissues across a range of barley developmental stages, phenotypic datasets from two consecutive years of field-grown trials in the United Kingdom, Germany and the USA; and whole genome shotgun sequencing from all cultivars, which was used to complement the RNA-sequencing data for variant calling. The outcomes are a filtered SNP marker file, a phenotypic database and a large gene expression dataset providing a comprehensive resource which allows for downstream analyses like genome wide association studies or expression associations.

## Background & Summary

Barley is one of the most important crops worldwide (5^th^ in 2020 on area harvested, FAOSTAT^1^) and has a high value in the European agricultural sector underpinning the beer and whisky industries^2^. New barley cultivars are introduced to the market every year, after being evaluated for multiple traits e.g., disease resistance, yield, and malting quality traits^3^. Barley breeding and the introduction of barley cultivars started at the beginning of the 19^th^ century in the UK and by the end of the 19^th^ century all over Europe^4^. Instead of seeds being grown by the farmer with some saved for subsequent sowing the following year, breeding institutes were established, with the mission to develop improved seed stocks. Early cultivars were developed through mass selection and later followed by line selection from landraces. Initial breeding efforts focused on increasing yield^5^. Due to the considerable success of these breeding efforts, seed stocks soon became distributed across the continent and each country started their own breeding program by incorporating local landraces in crosses with these generally higher yielding genotypes. This cross-breeding technique of simple crosses followed by selection quickly led to an increase in yield as shown for spring barley in Germany with a doubling of yield from 1800 to 1900^6^. Breeding developed further by intentionally mutating seeds with chemicals or radiation to induce higher genetic variation in the offspring^7^. One of the most notable results from mutation breeding were the dwarfing genes which were critical for the green revolution^8^. Shorter stature cultivars provided the advantage of preventing lodging which was crucial for the development of high-yielding cultivars with heavy spikes. Complementing traditional to cross- and mutation-breeding, molecular technologies developed further and were quickly adopted. One of the most successful advances was marker-assisted selection (MAS) which deploys molecular markers to detect allelic variations within a genome. The most common markers used in breeding nowadays are single nucleotide polymorphisms (SNPs)^9^. MAS is used for rapid and high-throughput selection of new genotypes and has matured from single marker analysis to genome-wide selection approaches. While SNPs are a key component of the genotyping platforms used in plant breeding purposes, they can also be used for gene discovery. Quantitative trait locus (QTL) mapping and genome wide association studies (GWAS) are valuable to identify alleles for genes underpinning genetically complex traits^10-13^. High throughput genetic markers are however only one of a number of genetic and genomic resources that have effectively revolutionised genetics and breeding. Next generation sequence data formed the basis of the first barley genome published in 2017 from the cultivar Morex^14^ which has been followed quickly by additional genomes from other cultivars^15^. The availability of “reference genome sequences” has both simplified the process and allowed a more precise identification of the causative genes controlling phenotypic traits.

Here we introduce new genetic and genomic datasets assembled from a European two-row spring barley population that is representative of pan-European breeding progress across the years from 1830 to 2014. A total of 209 50K SNP-array^16^ genotyped barley cultivars were selected and grown in replicated field trials across three contrasting environments and for two years to score agronomic traits. Six different tissues from each cultivar were harvested and RNA was isolated for the collection of tissue and genotype specific transcript abundance (RNA-seq) data. Using both this RNA-seq data and whole genome shotgun sequence data from all individuals in the population, an exhaustive collection of high confidence SNP markers was assembled. We describe these datasets and provide examples of how they can be used.

## Methods

### Barley material and field trials

We assembled a collection of 209 European two-rowed spring barley cultivars (Supplemental Table 1), which is a representative subset of previously described two-row spring European barley populations^10,11,17-19^ that show a significant increase in yield over time. Pedigree data was collected from publications^17,20^, and the following two websites: https://grinczech.vurv.cz/gringlobal/search.aspx and https://www.lfl.bayern.de/mam/cms07/ipz/dateien/abst_gerste.pdf. Field experiments were conducted at the Leibniz Institute of Plant Genetics and Crop Plant Research (**IPK**) in Gatersleben, Germany, the James Hutton Institute (**JHI**) in Dundee, UK and the University of Minnesota (**UMN**) in St. Paul, USA in 2019 and 2020. At IPK and UMN, 100 grains of each genotype were sown in 1 m long double-rows in a completely random design with three replications in both years. At JHI, a seed density estimated to produce 350 plants per m^2^ for plot sizes of 2m × 1.5 m was established. In 2019 a single replicate was grown and in 2020 a completely random design with two replicates. In addition, a polytunnel trial was included at JHI in 2019. Plant material was grown in 7 litre sized pots, 4 seeds per pot, in 3 replicate sets in a completely random design. Each replicate set had 8 columns and 30 rows and contained a replicate of each of the 209 genotypes.

### Phenotyping

In total, 29 phenotypes were recorded on a per-plot basis in the field trials or on a per-pot basis in the polytunnel experiment. Developmental traits, growth habit and plant height measurements were recorded in the trials as described in Table 1. To measure spike and grain traits, ten to 15 main tiller spikes were harvested at full maturity (Zadoks stage 92) per plot, excluding the outermost plants of each row to avoid edge effects. After recording of all spike traits, spikes were hand-threshed, and grains were subjected to size and weight measurements on a Marvin SeedAnalyzer 6 (MARViTECH GmbH, Germany). Samples were first weighted and then added on to the Marvin tray for optical measurements of the grain size.

**Table 1:** A summary of the phenotypic traits and a description on how they were scored.

| Trait | Method |
| --- | --- |
| **Developmental traits** |  |
| Days to awn tipping | 50% of the main tiller awns per plot have emerged up to 1 cm out of the flag leaf sheath. Recorded as days since sowing |
| Days to heading | 50% of the main tiller spikes per plot have emerged halfway out of the flag leaf sheath. Recorded as days since sowing |
| Days to senescence | 50% of the main tiller peduncles per plot are senescent (yellow). Recorded as days since sowing |
| Days from awn tipping to heading | Derived from days to awn tipping and days to heading |
| Days from awn tipping to senescence | Derived from days to awn tipping and days to senescence |
| Days from heading to senescence | Derived from days to heading and days to senescence |
| Growth habit (GH) | Visual evaluation using a scale of 1 (erect), 2 (intermediate) and 3 (prostrate). Recorded at the onset of stem elongation |
| **Height and length traits** |  |
| Peduncle base height | Height of the base of the peduncle in cm |
| Flag leaf blade height | Height of the flag leaf sheath in cm |
| Culm height | Height of the base of the spike in cm |
| Plant height | Height of the top of the spike in cm |
| Awn tip height | Height of the tip of the awns in cm |
| Spike base to flag leaf | Calculated distance from base of spike to flag leaf sheath (auricle) in cm |
| Peduncle length | Calculated distance from base of spike to base of peduncle in cm |
| Awn length | Calculated distance from tip of awns to top of spike in cm |
| Spike culm ratio | Spike length divided by culm height |
| **Spike traits (recorded on 10-15 main tiller traits per plot after harvest)** |  |
| Rachis node number | Number of rachis nodes |
| Spike length | Spike length in cm |
| Spike density | Rachis node number divided by spike length |
| **Grain traits** |  |
| **Grain traits (recorded on 10-15 main tiller traits per plot after harvest) using a Marvin Seed Analyzer 6** |  |
| Grain area | Area of all kernels per spike in mm<sup>2</sup>. Recorded using the automatic grain area calculation function in the Marvin SeedAnalyzer 6 software |
| Kernel roundness | Roundness of all kernels per spike. Recorded using the automatic kernel roundness calculation function in the Marvin SeedAnalyzer 6 software |
| Thousand kernel weight | Calculated from the number and weight of the kernels using the Marvin SeedAnalyzer 6 software |
| Grain length | Length of all kernels per spike in cm |
| Grain width | Width of all kernels per spike in cm |
| **Spike traits (recorded on 10-15 main tiller traits per plot after harvest)** |  |
| Infertile florets at top and bottom (=edges) of spike | Number of infertile florets at the top of spike down to first fertile floret + number of infertile florets at the base of spike up to first fertile floret |
| Infertile florets in the middle of the spike | Number of infertile florets in the centre of the spike |
| Number of fertile grain | Total number of fertile florets per spike |
| Percent of fertile florets | Total number of fertile florets per spike divided by rachis node number |
| Number infertile florets | Total number of infertile florets per spike |

### Tissue sampling for RNA-seq

Six different tissues were sampled for RNA-seq analysis: crown, root, inflorescence, peduncle, spikelet and grain. At UMN, crown and root tissues were sampled from seven-day-old seedlings (GRO:0007060, first leaf unfolded). Ten seeds per genotype were surface sterilized and planted in moist vermiculite in individual Cone-tainers (6000 RLC3 size, Ray Leach, Tangent, OR). The Cone-tainers were put into a dark cold room for four days to achieve more consistent germination. Then they were moved into a growth chamber at 20°C with 16 hours of light for seven days. Tissues were harvested within three hours, starting at 9:00 am USA Central Time Zone to reduce the circadian effect on gene expression. Roots were sampled by cutting the longest root from each seedling adjacent to the germinated seed, and crowns by removing the roots and keeping the 1 cm shoot tissue immediately above. For each individual genotype five plants were combined and snap frozen in liquid nitrogen.

At JHI, the barley plants grown in 2019 under polytunnel conditions were used for tissue sampling. When plants reached the booting stage, which was 84-85 days after germination, 3 – 5 cm whole developing inflorescence tissue was taken, two from each replicate per genotype per sample. Whole peduncles were taken at 2 – 5 cm in length, three from each replicate per genotype per sample when plants were 88 – 90 days old. Samples were snap frozen in liquid nitrogen and stored at -80°C.

At IPK, barley plants grown in the 2019 field trial were monitored daily by dissecting single spikelets and recording the date of green anther stage and flowering stage on paper tags attached to the spikes. Sampling was limited to a two-hour period between 10:00 and 12:00 am Central European Summer Time each day to reduce the circadian effect on gene expression. Three spikes per plot of one repetition were selected at green anther stage and two central spikelets from the centre of each spike were sampled. At 5 days post anthesis, three spikes per plot of one repetition were selected and six developing grains per spike were sampled from the central region of each spike. All samples were snap-frozen in liquid nitrogen and stored at -80 °C until RNA extraction.

### RNA extraction and RNA sequencing

RNA was extracted using the RNeasy Plant Mini Kit (Qiagen) and treated with DNaseI following the manufacturer’s instructions. Buffer RLC was used for seedling root extractions, and Buffer RLT was used for all other tissue extractions. To ensure a high purity of spikelet and grain samples, a more rigorous cleanup using 700µl RW1 and three wash steps with RPE was performed. The integrity of samples was determined using an Agilent 2100 Bioanalyzer, an Agilent 4200 TapeStation or a 1% agarose gel. All tested samples had a RIN factor of >=8 and were suitable for further processing. Paired-end libraries with a read length of 150 bp were constructed from spikelet and grain samples (IPK Gatersleben) and seedling root and crown samples (University of Minnesota Genomics Center, Minneapolis, MN, USA) using the Illumina TruSeq Stranded Total RNA Library Prep Plant with Ribo-Zero Plant kit and sequenced on the NovaSeq 6000 platform. For the inflorescence and peduncle samples (JHI) Illumina RNA-seq library preparation and RNA-seq was carried out by Novogene (Company Limited, Hong Kong). The libraries were prepared using NEBNext® Ultra™ Directional RNA Library Prep Kit and sequenced using Illumina NovaSeq 6000 (PE 150).

## Bioinformatics

### Read quantification

We generated 77.95 billion raw reads from RNA-seq of the six different tissues (Supplemental Table 2). Raw reads were trimmed with Trimmomatic 0.39^21^ to remove adapters and reads shorter than 60 bp. Salmon 1.3.0^22^ was used for expression quantification including the gcBias setting to align trimmed reads to the transcriptome. We followed the approach of selective alignments by generating a decoy-aware transcriptome from the barley reference transcript dataset V2 (BaRTv2)^23^ and the reference genome of **cv** Barke^15^. This approach is recommended^24^ to reduce inaccurate transcript quantification caused by unannotated genomic loci that have a high sequence similarity to annotated transcripts.

### Expression analysis

Tissue-specific genes were identified using different R packages^25^. For each tissue the raw counts were imported and combined to gene expression counts using tximport^26^. Raw counts were normalised (calcNormFactor), and log transformed to counts per million (cpm) using edgeR^27^. The tissue-specific expressed genes were identified by filtering for an average cpm of above 1 across all samples in this tissue and an average cpm of below -1 for all the other tissues. In addition, gene expression for two and more tissues were filtered with the same parameters to build the intersection sets required to create an UpSet^28,29^ plot of expressed genes in the different tissues. Gene ontology (GO) enrichment for the identified genes and visualisation was done as previously described^30^.

### Variant calling

For variant calling, the trimmed RNA-seq reads were mapped to the reference genome of **cv** Barke^15^ using the two-pass mode implemented in STAR v. 2.7.5^31^ allowing 6% mismatches normalized to read length, intron lengths between 60 and 15000 bp, a maximum distance of 2000 bp between mates and a maximum number of 30,000 transcripts per window. Due to the high number of reads in the grain and spikelet tissue the splice junction files were filtered for at least one uniquely-mapped read in more than one sample, with non-canonical splice sites removed and then used to generate a new genome index for the second mapping run. For the other tissues, the splice junction files from the first pass were provided as part of the input for the second mapping step. Duplicated reads were marked with Picard 2.18.29^32^ followed by filtering with bamtools 2.5.1^33^ to remove reads with 2% mismatches and a mapping quality <= 50. The legacy algorithm of Freebayes 1.3.2^34^ was used to call variants with a minimum fraction of alternate allele observations of 20%, a minimum alternate allele count of 2, a minimum coverage of 4, and minimum base and mapping qualities of 30.

### Whole genome shotgun (WGS) approach

DNA was extracted from snap-frozen second leaves of greenhouse-grown (21°C/18°C day/night temperature) two-week old seedlings using a guanidinium thiocyanate-NaCl-based method as described^35^. DNA quality and quantity was assessed by agarose gel electrophoresis. The Nextera DNA kit (Illumina) was used for library preparation of 150 bp paired-end reads. Libraries were multiplexed and sequenced on a NovaSeq 6000 platform at IPK Gatersleben. 12.16 billion raw paired-end raw reads (Supplemental Table 2) were trimmed with Cutadapt 1.15^36^ to remove adapters and reads shorter than 30 bp. Trimmed reads were mapped to the reference genome of **cv** Barke using Minimap2 2.11^37^. The resulting alignment files were sorted and duplicate-marked using Novosort 3.06.05^38^ and converted to cram files using samtools 1.8^39,40^. Bcftools^40^ call was used to call variants using genotype likelihoods calculated from alignments with a minimum quality score of 20 with bcftools mpileup. Variants were re-called based on read depth ratios using a custom awk script similar to the one at https://bitbucket.org/ipk_dg_public/vcf_filtering/src/master/ with the following parameters modified: dphom = 1, dphet = 2, minhomn = 10, tol = 0.249, minmaf = 0.1, minpresent = 0.01.

### Genotype marker file

The final genotype file was generated by filtering and merging multiple files. First all RNA-seq vcf files from the six tissues were filtered to remove insertions and deletions (Indels). RNAseq SNPs from individual tissues were then merged, prioritizing homozygous calls by retaining heterozygous calls only if no homozygous calls were present in any of the tissues. The merged RNAseq SNPs were then merged with WGS and 50K array SNPs^16^, prioritizing homozygous calls in the same manner. The resulting unfiltered dataset contained 209 cultivars and 32,484,981 bi-allelic SNPs. The merged SNP dataset was filtered using TASSEL5^41^ to remove SNPs with more than 20% missing data, minor allele frequency (MAF) < 0.01, heterozygosity > 0.02, and only keeping bi-allelic SNPs. Missing data was imputed using the FILLIN plugin^42^ in TASSEL5 by first identifying haplotypes. For haplotype identification each chromosome was split into 500 blocks. The number of markers per haplotype block (-hapSize) was the total number of markers per chromosome divided by 500 and rounded to be divisible by 64 (TASSEL5 software requirement). Haplotypes were identified for each block with a maximum number of haplotypes of 20 (-maxHap 20) and at least five different genotypes per haplotype (-minTaxa 5). Haplotype information was used as input for the imputation. Further filtering removed seven lines that had more than 30% missing data after imputation (Aramir, Balder J, Dallas, KWS Irina, Power, Proctor and Spey), and one line was removed that had more than 2% heterozygosity (Rika). In a last filtering step, we removed SNPs which still had more than 20% missing data, MAF < 0.025 and heterozygosity > 0.02. SNPs were LD pruned with PLINK (v1.9)^43^ using a window size of 5000, a step size of 50 and an r^2^ threshold of 0.99. The final SNP dataset after pruning contained 201 cultivars and 1,509,447 SNPs.

### Variant effect using SnpEff

To identify the effect of variants on the protein, we filtered the raw vcf files in a different way to generate an input file for SnpEff^44^. The aim for the genotype marker set explained above was to reduce the number of SNPs with pruning to a size which can be used for association analysis. For the variant effect, on the other hand, we need all the available SNP information and more importantly do not want to lose any SNPs due to pruning in gene space. For SNPs, the merged unfiltered vcf file containing RNA-seq, WGS and 50k data was filtered by removing heterozygous calls, removing SNPs with missing data in more than 20% of the samples and a minor allele frequency of < 0.025. In addition, a dataset containing Indels was created by using the six vcf output files from the RNA-seq data after variant calling with Freebayes. All were filtered to keep Indels only, remove heterozygous calls, removing variants with missing data in more than 20% of the samples and a MAF of < 0.025.

The six Indel vcf files were combined into one. A SnpEff database was built based on BaRTv2 and the Barke reference genome.

Statistical analysis of phenotypic data, calculation of Best linear unbiased predictions (BLUPs) and heritability Statistical analysis was performed using R 3.6.1^25^. The Pearson correlation coefficient between experiments was calculated for each phenotypic trait and datasets showing an insignificant correlation (p>0.05) with at least one other dataset of the same trait were removed before calculating BLUPs. The datasets being used in each of the BLUP calculations are listed in Supplemental Table 3. BLUPs were calculated across experiments using a randomized complete block model in META-R with experiments set as a random factor following formula 3 in Alvarado *et al*.^*45*^.

### Genome wide association studies (GWAS)

Association between phenotype and genotype was done using the Mixed Linear Model (MLM)^46^ with GAPIT (version 3)^47^. As input, we used the genotype marker file of 201 cultivars and 1,509,447 SNPs and the BLUP values of the Awn length phenotype were used to show an example of the process. Three principal components (PCs) were calculated within GAPIT and model selection set to TRUE to enable GAPIT to select the optimal number of PCs for the individual phenotype based on a Bayesian information criterion (BIC).

## Data Records

All raw data files for both raw RNA sequencing data and whole genome shotgun data have been deposited at the European Nucleotide Archive (ENA) under the following project number: PRJEB49069 for the RNA-sequencing reads and PRJEB48903 for the whole genome shotgun sequencing reads.

Phenotypic data and the SNP marker file are available through Germinate^48^: https://ics.hutton.ac.uk/germinate-barn/

The database contains the raw data by year and by site plus the calculated BLUP dataset.

Derived datasets are available through e!Dal^49^. The datasets consist of two sets of gene expression files per tissue; one for the raw read counts and one for the TPM values mapped against BaRTv2. Further two files containing the variant identification are available. The first contains the SNPs and the second with the Indel information with SnpEff annotation. Preview link: https://doi.ipk-gatersleben.de/DOI/815787d8-4036-408b-999e-725f7645eacf/6e532074-b4d7-4393-9221-8fe643e100f2/2/1847940088

## Technical Validation

### Population

The 209 two-row spring barley population was selected from previously established datasets containing 647 genotypes^10,11,17-19^. To include a wide range of genetically representative individuals, we used available BOPA SNP data (as previously described)^50^ and did a multi-dimensional scaling plot (Figure 1 a). Dimension 1 showed the progression from the oldest to the newest cultivars and explains 10.7% of the genetic variation. Genotypes were then chosen to be spread across year of registration as a cultivar to the UK market. Except for the first time-range which encompassed 130 years (1830-1959) of cultivar releases, all other ranges were split into decades and each time-range is represented by a similar number of genotypes (Figure 1 b). The final population was representative of breeding progress in cultivated barley for improved yield over time.

**Figure 1:**
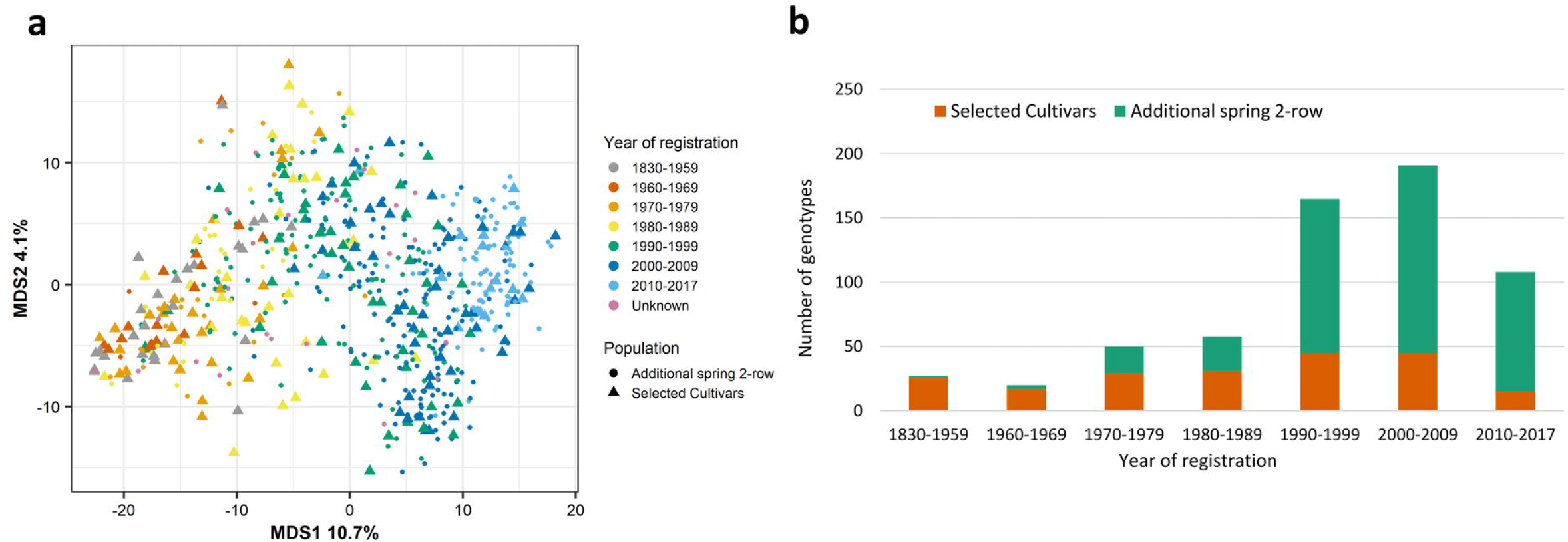
Selection of a two-row spring population. a) Multidimensional scaling plot of 647 European two-row spring cultivars. Genotype information came from 2,336 previously published BOPA markers. The 209 selected cultivars forming the population in this paper are shown as triangles. The year range represents the year each individual cultivar was registered. b) Distribution of the selected population of 209 cultivars (orange) as part of the total 647 European two-row spring cultivars (green) by year of registration.

Pedigree data showed that modern barley germplasm is highly connected with most current genotypes’ descendants of a small number of “founder” genotypes (Pedigree file: Supplemental File 1, Pedigree attributes: Supplemental File 2). These supplemental pedigree files can be used as input for the pedigree visualisation tool Helium (https://helium.hutton.ac.uk/)^51^. Intermediate crosses were omitted from the file to be able to display the pedigree and produce a tree which is both readable and navigable. Using the pedigree data within Helium allows for further analyses. For example, eigenvector centrality analysis can be used to measure the importance of a node in a network. The cultivar with the highest influence on the pedigree tree was Kenia, followed by Maja and Gull. In total, 163 out of 209 (77.6%) of the cultivars used in our population can be traced back to those three genotypes, confirming the importance of the founder genotypes for the overall population.

### Phenotyping

Field trials were done in 2019 and 2020 in three different locations: Minneapolis (Lat. 44.987, Long: -93.258; MN, USA), Dundee (Lat. 56.462, Long. -2.971; UK) and Gatersleben (Lat. 51.823, Long. 11.287; Germany). In total 29 agronomic traits were scored associated with development (earliness and growth habit traits), grain and height measurements. All the phenotypic data can be viewed and studied in a Germinate database: https://ics.hutton.ac.uk/germinate-barn/. Across years and sites, the results were consistent except for a few traits. All earliness traits showed a faster development to awn tipping all the way to peduncle senescence in Minnesota and slowest in Dundee, with for example the days from sowing to senescence differing by more than 50 days. One obvious outlier in the phenotypic scoring were the grain fertility measurements in Minnesota in 2019 where the spikes got stuck in the flag leaf sheath due to high temperatures during the growing season and did not emerge fully which led to a higher number of infertile florets. All phenotypic information was combined into BLUPs except for 24 out of 165 phenotype datasets which did not correlate with the rest (Supplemental Table 3 shows which phenotype values were combined; Supplemental Figure 1 for distribution of BLUP values per phenotype). The strongest positive correlation of the different phenotypes were the five different height measurements to each other (Figure 3). Those were also strongly negative correlated with year of registration. The earliness traits on the other hand showed a stronger positive correlation with year of registration (awn tipping to heading; awn tipping to senescence). Of the grain phenotypes, grain length and grain area were strongly correlated to each other and to thousand kernel weight, but less so to grain width. Interestingly, grain length is positively correlated with grain circularity, while grain width is negatively correlated with it.

### Genotyping

To achieve the most extensive genotypic information for our population, variant calling from RNA-seq data, whole genome shotgun data and previously established 50K SNP data was combined (32,484,981 raw SNPs). For RNA-seq and WGS, the data was filtered to keep only biallelic SNPs. We extracted the SNPs corresponding to the previous described BOPA markers across all 1463 sequencing datasets (six tissue-specific RNA-seq datasets with 209 genotypes each and one WGS dataset with 209 genotypes) for quality control. The Pearson correlation coefficient for all genotypes between datasets was calculated. This identified mixed-up samples where the genotype showed high correlation with a differently named sample and therefore allowed for correction of the genotypic information. Samples with a high number of heterogenous SNPs (above 10%) were removed as this pointed towards issues during sample preparation. The filtering step reduced the number of genotypes per tissue. The final numbers of genotypes per tissue varied between 191 to 199 (Supplemental Table 2 shows which genotypes per tissue where retained). The merged SNP file was filtered to remove highly heterozygous sites or those containing more than 20% missing data. The remaining sites were imputed using haplotype imputation. SNPs were pruned by LD using Plink to reduce the dataset size to the final 1,509,447 SNPs. SNP distribution along the 7 chromosomes showed overall low polymorphism in the centromeric region and higher polymorphism towards the telomeric ends (Figure 2). For chromosome 1H, 2H and 7H almost no polymorphism could be identified in the centromeric region, as previously observed by Mascher *et al* ^14^.

**Figure 2:**
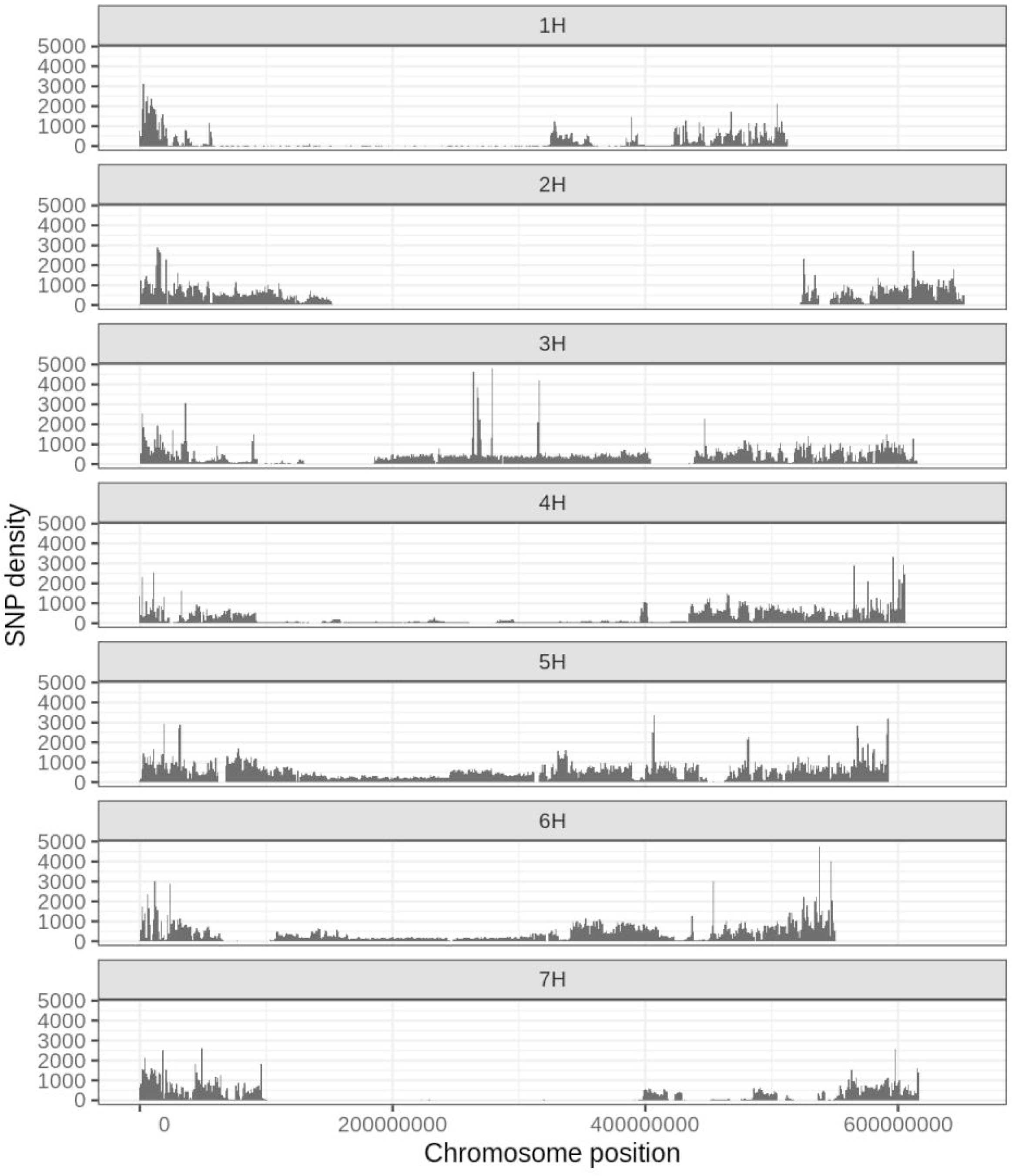
SNP distribution and density of the final 1,509,447 SNPs in the genotypic marker file along the seven barley chromosomes in 1 Mb bins. The SNPs were identified from the 50k SNP array, RNA-sequencing and whole genome shotgun sequencing datasets and filtered to remove missing values and heterozygosity.

**Figure 3:**
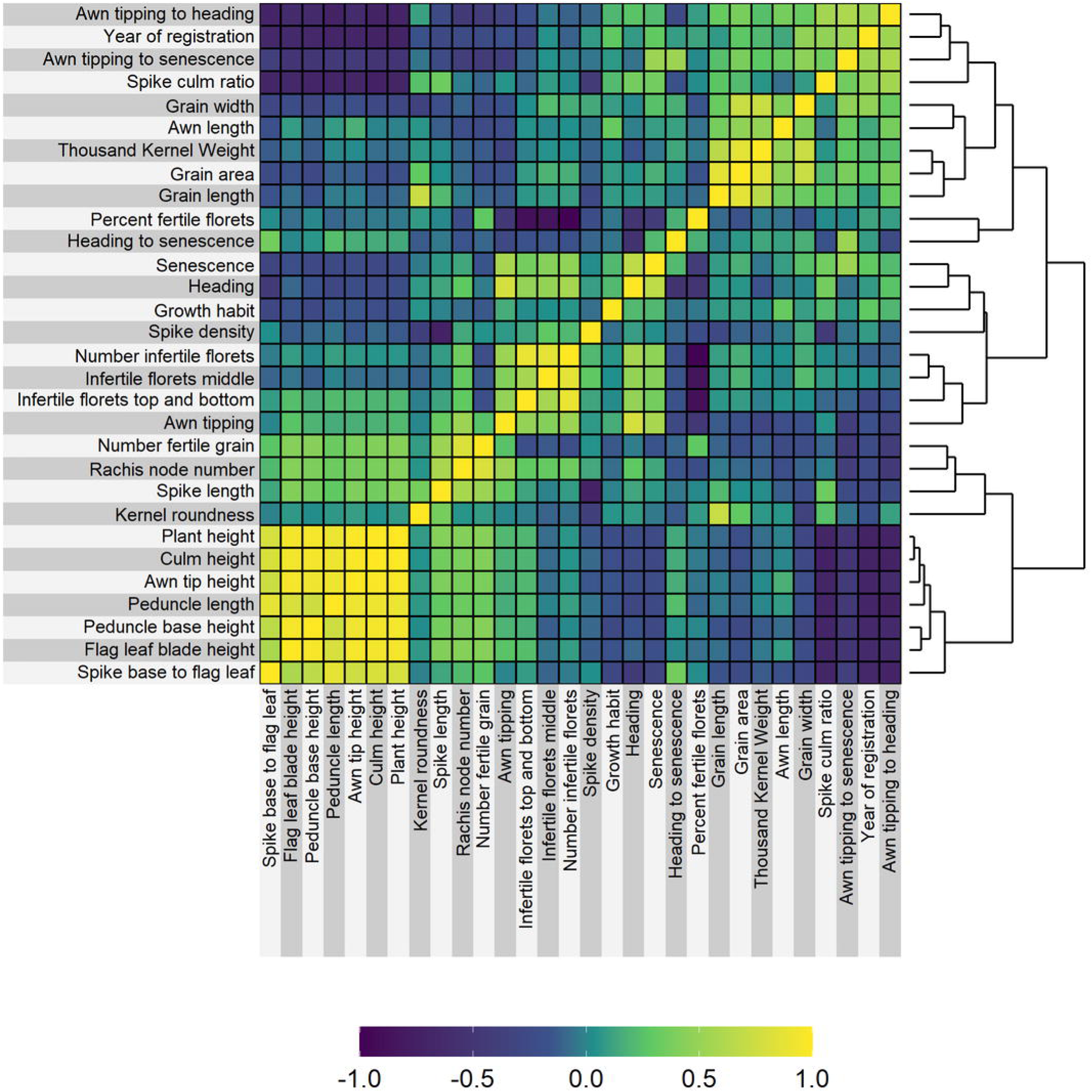
Pearson correlation coefficient between the 29 scored phenotypes and as 30th variable the year of registration. Phenotypic values were provided as best linear unbiased predictions for each phenotype for each of the 209 cultivars.

### Gene expression

RNA-seq data for six different tissues (crown, grain, inflorescence, peduncle, root, spikelet) was mapped against the BaRTv2 transcriptome using Salmon^22^. The expression of all 39,434 genes in transcript per million (TPM) for each tissue were used as input to generate a multidimensional scaling plot (MDS). The MDS shows all 209 genotypes cluster together by tissue type (Figure 4). The tissue furthest separated by the first dimension from the rest was the root tissue. The two tissues sampled from the spikelet at green anther stages (spikelet) and developing grain at five days post anthesis (grain) show the highest overlap.

**Figure 4:**
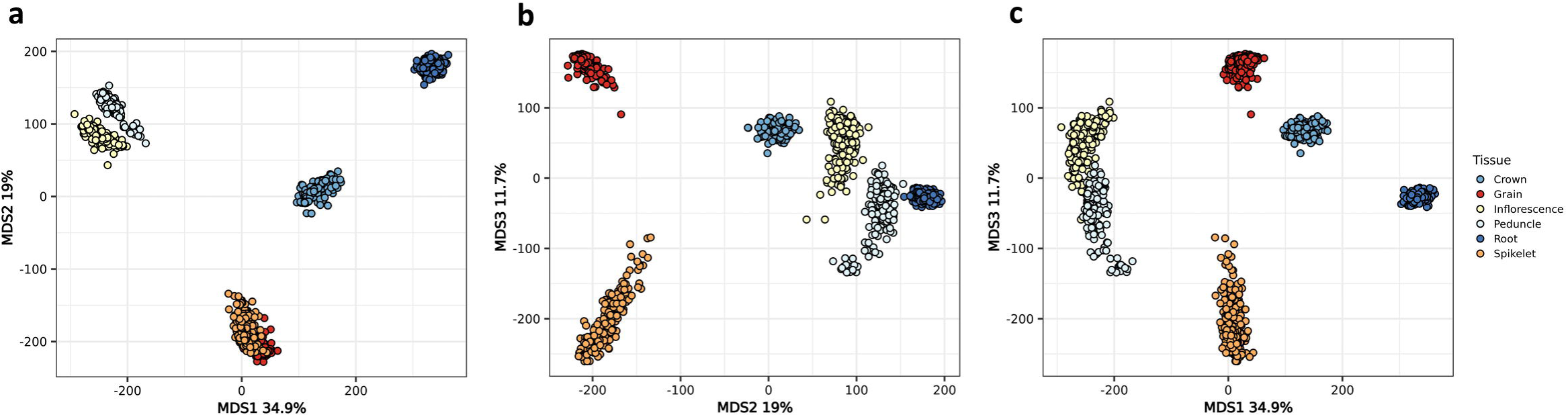
Gene expression of 209 cultivars across six tissues. Multidimensional scaling plot of all genes expressed in any of the six studied tissues: root, crown, peduncle, inflorescence, spikelet and grain.

### Data use-case scenarios

In the following three examples we show how the above datasets can be used.

In the first example the expression data has been used to filter for tissue-specific gene expression. Tissue-specific genes showed that the root tissue was the most distinct with 776 genes identified as root specific (Figure 5). 572 genes were specifically expressed in grain, 437 in spikelet, 198 in inflorescence, 86 in peduncle and 64 in crown. Inflorescence and peduncle shared the highest overlap of expressed genes with 927 genes and 13,215 genes were expressed in all six tissues. While the MDS plot shows a high overlap of samples between spikelet and grain in the first two dimensions, the third dimension divides those tissues which fits with these two tissues showing the second and third highest tissue-specific gene expression. Gene ontology for the peduncle resulted in no significant terms. The Gene ontology results for all remaining five tissues are shown in Figure 6. Most terms were identified for the root tissue showing terms one might expect in roots. Nitrate transport and general transmembrane transport were identified in the Biological Process category; apoplast, cell wall and extracellular region for the Cellular Compartment category. Different transferase activities are highly significant in the Molecular Function category as well as heme binding. The associated terms compare to those previously identified in maize^52^.

**Figure 5:**
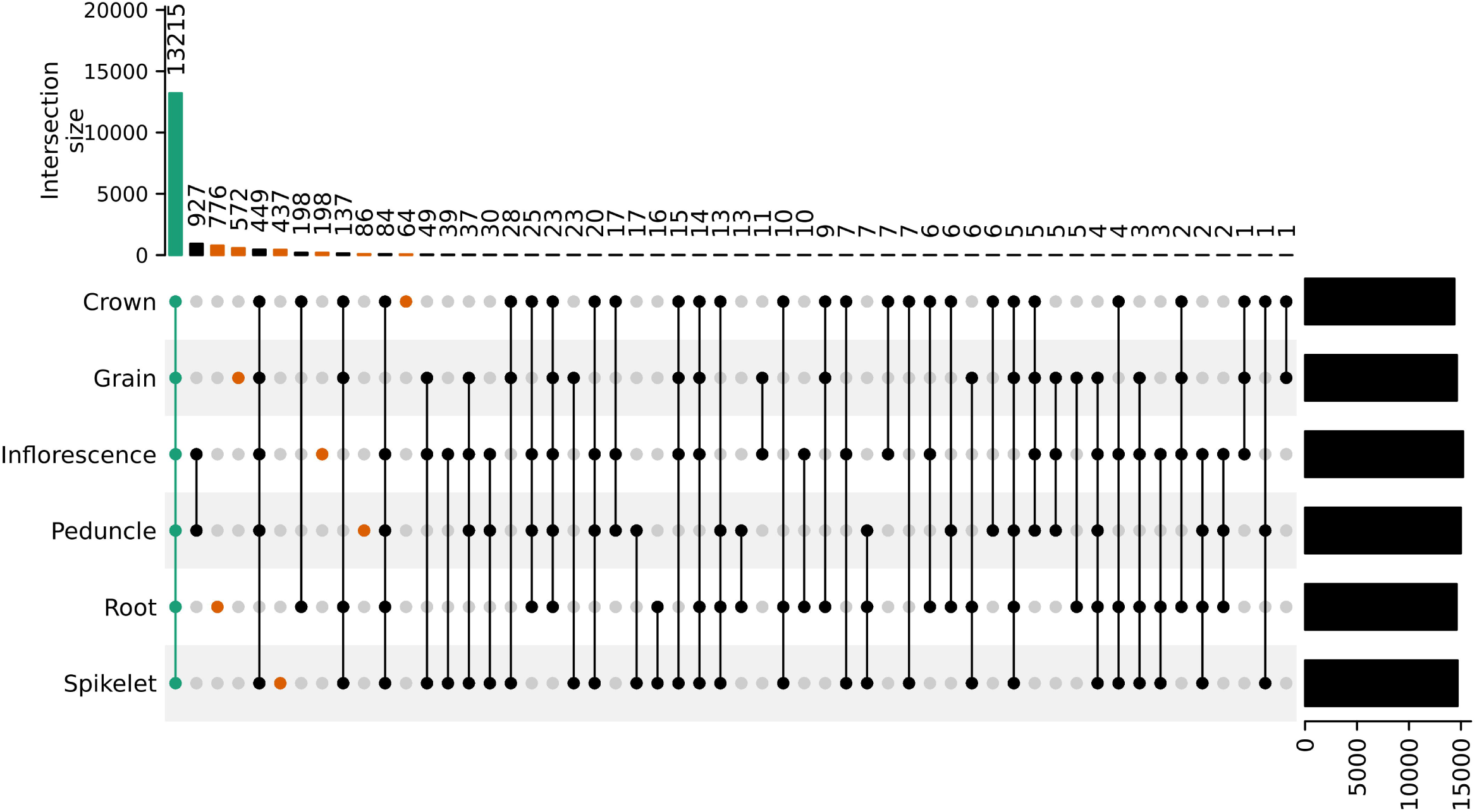
An UpSet plot showing the overlap of the expressed genes for each of the tissues and tissue combinations.

**Figure 6:**
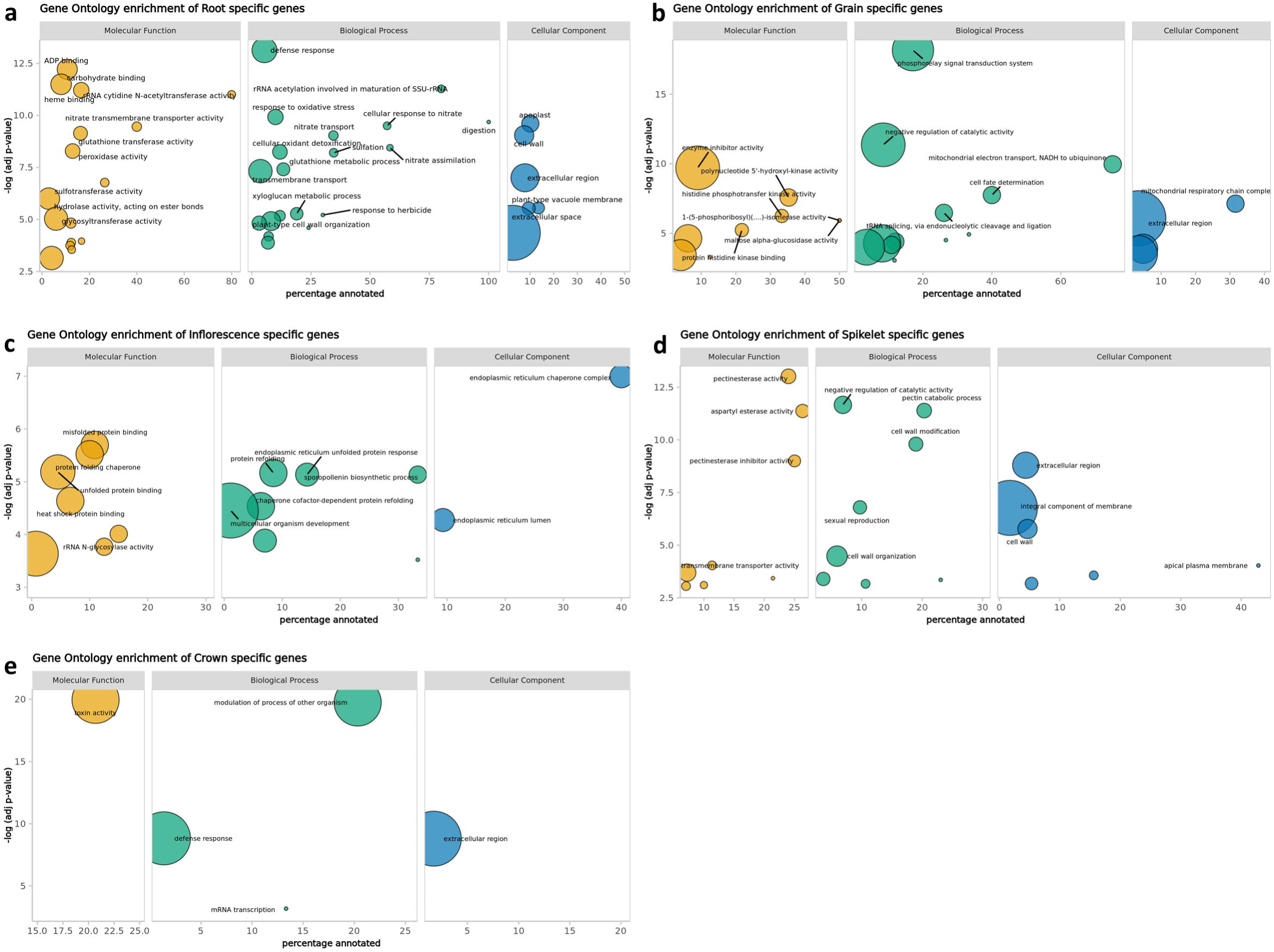
Gene ontology enrichment for the tissue-specific genes a) root, b) grain, c) inflorescence, d) spikelet and e) crown. X-axis shows the percentage of genes associated with the GO term out of all genes in BaRTv2 associated with this term. Y-axis shows the significance as FDR adjusted -log(p-value) of the GO term. The area of the circle corresponds to the number of genes associated with the GO term.

In the second example, we studied the potential impact that genetic variation could have on the gene activity or protein function by identifying premature stop codons or frameshift mutations in a high confidence variant dataset. For the SNP dataset we started with 32 million SNPs, removed heterozygous SNPs and filtered for variants with less than 20% missing data and a minor allele frequency of 2.5% which resulted in 4,012,229 SNPs^53^. Those were used as input into SnpEff which identified 9,219,271 effects (as described by SnpEff: http://pcingola.github.io/SnpEff/se_inputoutput/#eff-field-vcf-output-files) caused by those 4 million SNPs. 4% (368,650) of the effects were in exons, with 53.78% synonymous variants, corresponding to 199,545 effects in 17,446 genes. 45.57% (169,105 effects) of the exon effects were non-synonymous variants in 19,057 genes and 0.65% classified as nonsense. The 0.65% corresponded to 2,425 transcripts and 1,105 genes with a premature stop codon in the sequence. For the Indel identification only the RNA-seq variant files were considered as those provided higher read depth for the genetic regions. They were also filtered by removing heterozygous variants, keeping those with less than 20% missing data and a minor allele frequency of 2.5%. A total of 50,865 variants remained which SnpEff predicted to cause 558,991 effects. 50.31% (281,228 effects) were upstream or downstream of the gene and 41.81% (233,706 effects) in the intronic region. After filtering for disruptive frame shifts caused by insertions or deletions resulting in changes to the protein sequence, 1,912 genes remained which we designated as potentially non-functional in some of the cultivars^53^. Such structural variation can of course be explored in relation to gene expression. To illustrate, within the tissue-specific gene expression data we highlighted two genes (BaRT2v18chr5HG260690 and BaRT2v18chr2HG058650) with frameshift mutations in comparison to the Barke reference allele. Figure 7 shows the expression of those two genes. For BaRT2v18chr5HG260690 the deletion of four nucleotides is associated with lower gene expression. For BaRT2v18chr2HG058650 an A insertion is sociated with higher gene expression. The consequence of such variation still needs to be explored.

**Figure 7:**
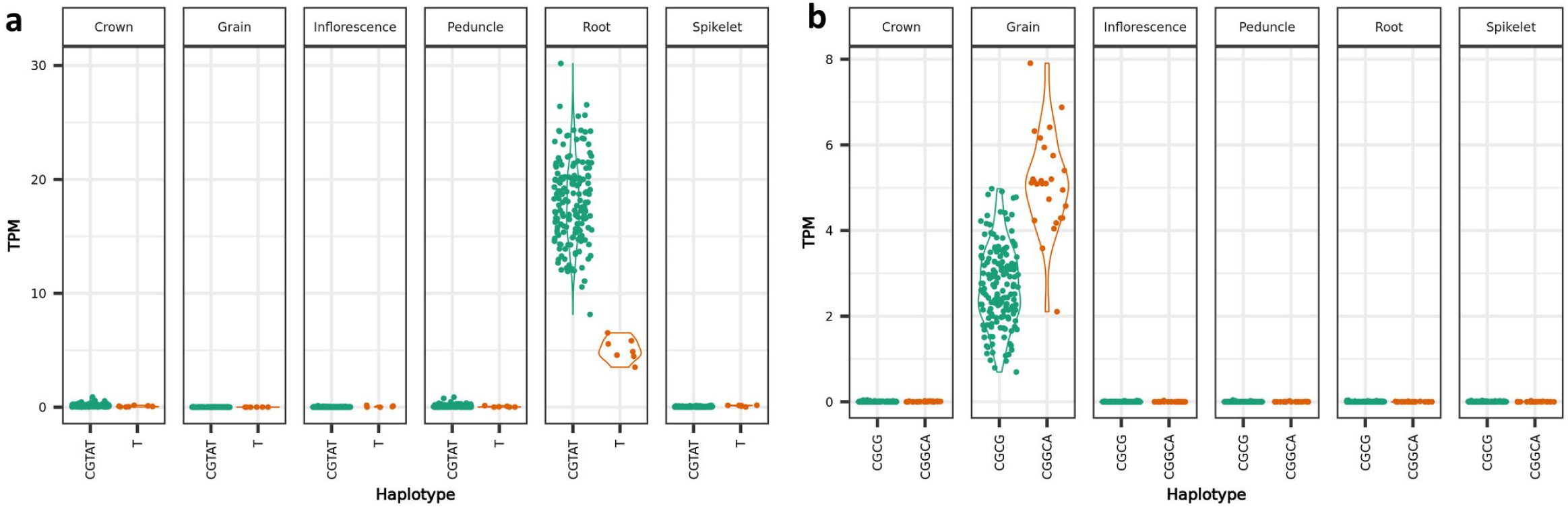
Gene expression in TPM (transcripts per million) for two genes identified with changes in the protein sequence. a) BaRT2v18chr5HG260690 and b) BaRT2v18chr2HG058650 split by haplotype on the x-axis. The first haplotype always represents the reference allele from the genotype Barke, and the second allele represents the alternative.

As a final example, we show a genome wide association study (GWAS) using the 1,509,447 SNP markers and the awn length phenotype. We used the Mixed Linear Model (MLM) in Gapit^47^ to identify associations in the genome. Using a -log10(p) cut-off of 5 resulted in 6 significant peaks (Figure 8). The most significant SNP was found on chromosome 5H at position 441Mb. This peak had a very rare MAF of 3% and was located within 1kb of a previously identified gene involved in awn length, *HvDep1* (BaRT2v18chr5HG247460). The here identified allele corresponded to the previously described ari-eGP allele on chromosome 5H which is a one base pair “A” insertion in the *HvDep1* gene^54^. The insertion was confirmed for four cultivars in the population: Golden Promise, Tyne, Midas and Chad. Gene expression of *HvDep1* (Figure 8b) showed potentially lower expression in crown and inflorescence tissue but similar levels for the peduncle. This simple example highlights the fact that variants causing a loss of function do not necessarily induce strong changes in gene expression.

**Figure 8:**
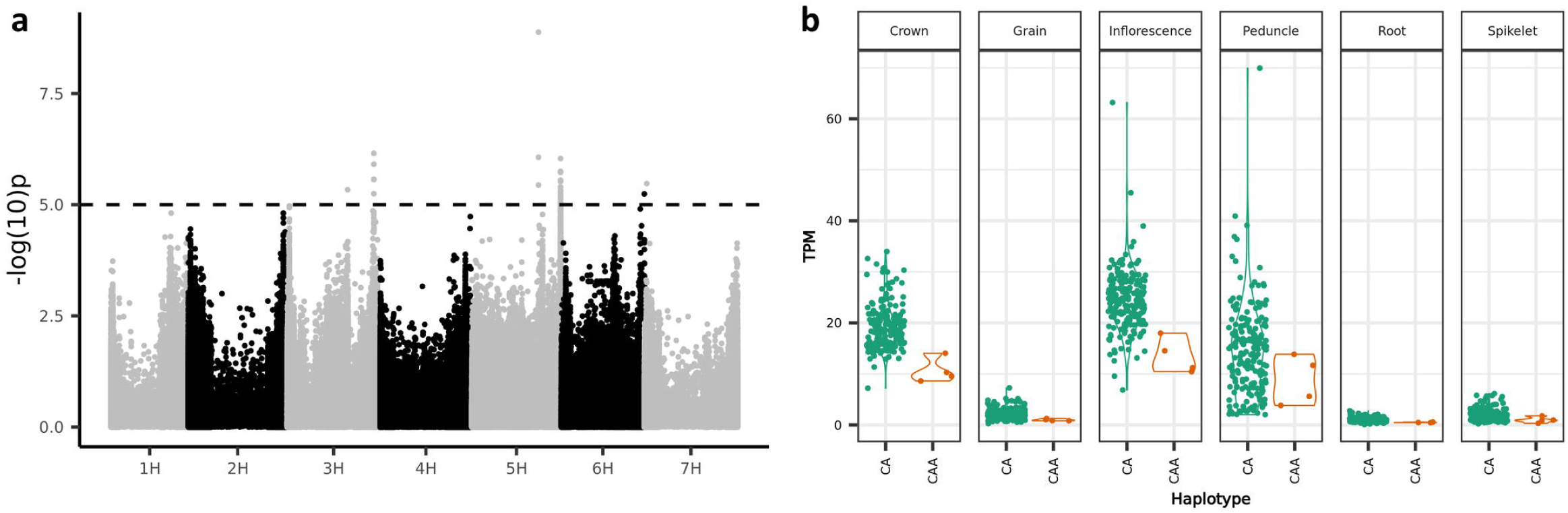
Genome wide association of awn length. a) GWAS result of awn length showing a high significant association on chromosome 5H. b) The gene expression in TPM of the candidate gene *HvDep1* (BaRT2v18chr5HG247460). The first haplotype on the x-axis represents the reference allele from the genotype Barke, and the second allele represents the alternative.

## Usage Notes

To perform the analysis using the Snakemake^55^ pipeline (see code availability) a high-performance computing (HPC) cluster is needed. For example, the Salmon indexing step in this setup needed 56Gb of memory using 16 cores, mapping of each individual sample needed 31Gb of memory using 8 cores. Downstream analyses like the genome wide association studies can be performed by downloading the BLUPs of the phenotypes and the marker file from Germinate.

## Code Availability

The code for analysing the RNA-sequencing data from mapping to genome and transcriptome to variant calling was combined into a Snakemake^55^ pipeline and is available on GitHub: https://github.com/SchreiberM/BARN.

## Acknowledgements

The authors would like to acknowledge Richard Keith and Chris Warden for maintaining the JHI field trials, Nicola McCallum and Ruth Hamilton for the help with phenotypic scoring and John Fuller for the RNA-seq library preparation. We thank Micha Bayer, Runxuan Zhang, John Brown and Wenbin Guo for advice on the project design and bioinformatics. We would like to acknowledge Shane Heinen, Yadong Huang, Ismaél Pfeifer, and Letícia Pasqualino for the help in the lab and field work at the University of Minnesota. We acknowledge current and former members of the Genomics of Genetic Resources research group at IPK Gatersleben: Mary Ziems for planning and maintaining field trials, phenotypic scoring and tissue sampling; Mark Timothy Rabanus-Wallace, Hélène Pidon, Sudharsan Padmarasu, Mingjiu Li, Jayavardhan Reddy Kunam, Mohammed Rafaqat, Mohammad Awais, Jaqueline Pohl, Susanne König, Ines Walde, Manuela Knauft, Manuela Kretschmann and Beate Kamm for phenotypic scoring and tissue sampling. We thank Ines Walde and Susanne König at IPK for preparing sequencing libraries and generating sequencing data.

Thanks are also given to the Research/Scientific Computing teams at The James Hutton Institute and NIAB for providing computational resources and technical support for the “UK’s Crop Diversity Bioinformatics HPC” (BBSRC grant BB/S019669/1), use of which has contributed to the results reported. The work was supported by funding from the Rural and Environment Science and Analytical Services Division of the Scottish Government. The authors acknowledge the Minnesota Supercomputing Institute at the University of Minnesota.

The project received funding in frame of the ERA-CAPS Research Programme with funding (i) of the German partners through German Research Foundation (DFG) to NS under the project references MU 3589/1-1 | STE 1102/15-1 | WA 3336/4-1, (ii) the Scottish partner to RW through BBSRC (award number BB/S004610/1), (iii) the US partner to GJM through NSF (award number 1844331)

## Author contributions

MS wrote the manuscript, conducted data collection and analysis. RWo conducted RNA-seq and WGS, data collection and analysis. AHa conducted RNA-seq data collection and analysis. MC did RNA-seq and data collection. JR provided seed material and did field trials. AHi conducted RNA sequencing, whole genome shotgun sequencing and primary data analysis. AF did data management. GJM, NS and RWa designed the experiment and supervised the work.

## Competing interests

There is no conflict of interest.

Supplemental Figure 1: Violin and boxplots of BLUP values for the phenotypic variation in all 29 scored traits.

