## Supplemental Table 1 for "Genomic resources for a historical collection of cultivated two-row European spring barley genotypes"

| Genotype | Country of Breeding Institute | Year of registration | Genotype | Country of Breeding Institute | Year of registration |
| --- | --- | --- | --- | --- | --- |
| Aapo | FIN | 1975 | Krystal | CZE | 1981 |
| Abacus | GBR | 1974 | KWS Irina | GBR | 2011 |
| Abava | LVA | 1978 | KWS Orphelia | GBR | 2010 |
| Acoustic | GBR | 2011 | KWS Vitara | GBR | 2014 |
| Alabama | DEU | 1998 | Kym | GBR | 1979 |
| Alexis | DEU | 1986 | Landlord | GBR | 1994 |
| Alliot | DNK | 1998 | Lenta | DNK | 1943 |
| Aluminium | DNK | 2003 | Livet | GBR | 1996 |
| Alva | SWE | 1977 | Lud | GBR | 1976 |
| Anais | DEU | 1999 | Macaw | FRA | 2002 |
| Annabell | DEU | 1997 | Maja | DNK | 1927 |
| Apex | NLD | 1983 | Mala | DNK | 1972 |
| Appaloosa | GBR | 2003 | Maresi | DEU | 1986 |
| Aramir | NLD | 1972 | Maris Mink | GBR | 1973 |
| Armelle | FRA | 1974 | Maypole | GBR | 2000 |
| Arvo | FIN | 1966 | Meltan | SWE | 1990 |
| Athos | FRA | 1975 | Midas | GBR | 1970 |
| Atlas | CZE | 1976 | Momentum | GBR | 2012 |
| Avalon | DEU | 2011 | Natasha | FRA | 1982 |
| Avec | SWE | 1995 | Natasia | GBR | 2010 |
| Balder | SWE | 1942 | Nomad | GBR | 1987 |
| Balder J | SWE | 1964 | Nordal | DNK | 1971 |
| Balga | LVA | 1990 | Novello | GBR | 2000 |
| Barke | DEU | 1996 | Odyssey | FRA | 2009 |
| Baronesse | DEU | 1989 | Okos | ITA | 1975 |
| Beatrix | DEU | 2003 | Olympus |  | 2013 |
| Beka | FRA | 1954 | Optic | GBR | 1992 |
| Berac | NLD | 1969 | Orbit | SVK | 1986 |
| Berenice | FRA | 1972 | Otto | ITA | 1972 |
| Berwick | GBR | 1997 | Overture | FRA | 2009 |
| Betzes | DEU | 1957 | Paloma | DNK | 1996 |
| Binder | DNK | 1913 | Pewter | GBR | 1998 |
| Binder Abed | DNK | 1913 | Pitcher | GBR | 1993 |
| Birgitta | SWE | 1963 | Poker | GBR | 2003 |
| Blenheim | GBR | 1984 | Potter | SWE | 1997 |
| Bogart | GBR | 2009 | Power | DNK | 2002 |
| Bonus | SWE | 1950 | Prague | GBR | 2004 |
| Braemar | GBR | 1999 | Prestige | GBR | 1998 |
| Brazil | FRA | 2000 | Prisma | NLD | 1985 |
| Britta | SWE | 1964 | Proctor | GBR | 1952 |
| Camargue | DEU | 1983 | Prodigal | GBR | 2010 |
| Campala | FRA | 2000 | Propino | GBR | 2007 |
| Carlsberg | DNK | 1946 | Publican | GBR | 2004 |
| Cellar | GBR | 1998 | Quench | GBR | 2004 |
| Centurion | GBR | 1996 | Rainbow | GBR | 1994 |
| Century | GBR | 1996 | Rapid | CZE | 1976 |
| Chad | GBR | 1988 | Reggae | NLD | 1992 |
| Chalice | GBR | 1995 | Renaissance |  | 2011 |
| Chariot | GBR | 1989 | Renata | GBR | 1994 |
| Chaser | GBR | 1997 | RGT Conquest | GBR | 2013 |
| Chevalier Tystofte 2 | GBR | 1830 | RGT Planet | GBR | 2014 |
| Chieftain | GBR | 1993 | Rhynchostar | GBR | 2010 |
| Chime | GBR | 1997 | Ria | DEU | 1987 |
| Cilla | SWE | 1966 | Rika | SWE | 1949 |
| Claret | GBR | 1980 | Riviera | GBR | 1992 |
| Class | GBR | 2000 | Romi | DNK | 1983 |
| Cocktail | GBR | 2003 | Rummy | GBR | 2003 |
| Colada | GBR | 1997 | Salka | DNK | 1973 |
| Concerto | GBR | 2006 | Saloon | GBR | 1997 |
| Cooper | GBR | 1991 | Salve | SWE | 1974 |
| Cork | GBR | 1992 | Scandium | DNK | 2004 |
| Corniche | DEU | 1983 | Scarlett | DEU | 1995 |
| Crusader | NLD | 1995 | Sebastian | DNK | 2000 |
| Dallas | GBR | 1989 | Senat | SWE | 1974 |
| Dandy | GBR | 1984 | Simba | DNK | 2003 |
| Deba Abed | DNK | 1965 | Simon | SWE | 1978 |
| Decanter | GBR | 1996 | Skittle | GBR | 2003 |
| Delibes | GBR | 1991 | Spartan | CZE | 1977 |
| Delta | NLD | 1959 | Spey | GBR | 1995 |
| Derkado | DEU | 1987 | Spire | GBR | 1999 |
| Diamant | CZE | 1965 | Starlight | GBR | 1997 |
| Digger | GBR | 1983 | Static | GBR | 1996 |
| Dina | DNK | 1975 | Steffi | DEU | 1989 |
| Domen | NOR | 1952 | Steina | DEU | 1980 |
| Drake | NLD | 1968 | Stendes | DNK | 1972 |
| Drost | DNK | 1954 | Sultan | NLD | 1966 |
| Drum | GBR | 2001 | Summit | GBR | 2008 |
| Egmont | GBR | 1980 | SW Scania | SWE | 2002 |
| Emir | NLD | 1962 | SY Taberna | GBR | 2008 |
| Fairytale | DNK | 2005 | SY Universal | GBR | 2009 |
| Favorit | CZE | 1973 | Tankard | GBR | 1993 |
| Forensic | GBR | 2006 | Taphouse | GBR | 2004 |
| Freja | SWE | 1941 | Tartan | GBR | 2004 |
| Georgie | GBR | 1973 | Tavern | GBR | 1997 |
| Gitane | NLD | 1976 | Tennis | GBR | 1984 |
| Golden Promise | GBR | 1966 | Tocada | DEU | 2002 |
| Golf | GBR | 1980 | Tremois | FRA | 1989 |
| Gull | SWE | 1913 | Trinity | GBR | 1993 |
| Gundel | SWE | 1984 | Triumph | DEU | 1973 |
| Hart | GBR | 1986 | Troubadour | FRA | 1982 |
| Hellas | SWE | 1967 | Tweed | GBR | 1980 |
| Helmi | FIN | 1942 | Tyne | GBR | 1985 |
| Hemingway | GBR | 2012 | Union | DEU | 1955 |
| Heron | GBR | 1990 | Vada | NLD | 1958 |
| Husky AFP2429 | GBR | 2006 | Valticky | CZE | 1950 |
| Imidis | GBR | 2006 | Vankkuri | FIN | 1943 |
| Impala | NLD | 1964 | Velvet |  | 2001 |
| Infinium |  | 2014 | Villa | DEU | 1968 |
| Ingrid | SWE | 1956 | Volla | DEU | 1957 |
| Invictus |  | 2013 | Waggon | GBR | 2002 |
| Isabella | DNK | 2004 | Westminster | GBR | 2002 |
| Isaria | DEU | 1924 | Wing | SWE | 1968 |
| Kassima | NLD | 2003 | Wisa | DEU | 1951 |
| Klaxon | GBR | 1981 | Zephyr 2RSB | NLD | 1965 |
| Koral | CZE | 1980 |  |  |  |
