## Supplementary figures and images for "Genomic resources for a historical collection of cultivated two-row European spring barley genotypes"

### Supplemental Figure 1

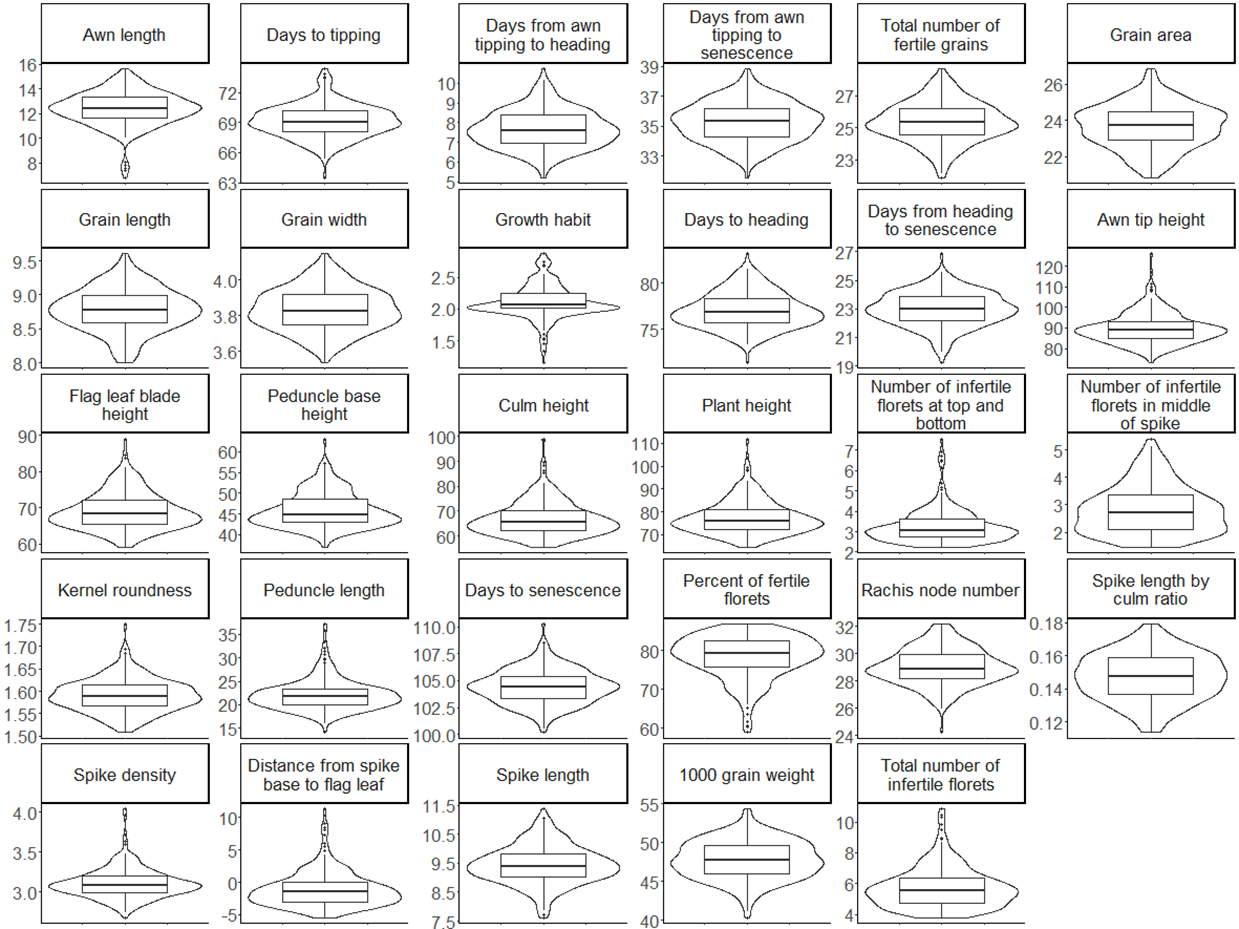
